# Limits to anatomical accuracy of diffusion tractography using modern approaches

**DOI:** 10.1101/392571

**Authors:** Kurt G Schilling, Vishwesh Nath, Colin Hansen, Prasanna Parvathaneni, Justin Blaber, Yurui Gao, Peter Neher, Dogu Baran Aydogan, Yonggang Shi, Mario Ocampo-Pineda, Simona Schiavi, Alessandro Daducci, Gabriel Girard, Muhamed Barakovic, Jonathan Rafael-Patino, David Romascano, Gaёtan Rensonnet, Marco Pizzolato, Alice Bates, Elda Fischi, Jean-Philippe Thiran, Erick J. Canales-Rodríguez, Chao Huang, Hongtu Zhu, Liming Zhong, Ryan Cabeen, Arthur W Toga, Francois Rheault, Guillaume Theaud, Jean-Christophe Houde, Jasmeen Sidhu, Maxime Chamberland, Carl-Fredrik Westin, Tim B. Dyrby, Ragini Verma, Yogesh Rathi, M Okan Irfanoglu, Cibu Thomas, Carlo Pierpaoli, Maxime Descoteaux, Adam W Anderson, Bennett A Landman

## Abstract

Diffusion MRI fiber tractography is widely used to probe the structural connectivity of thebrain, with a range of applications in both clinical and basic neuroscience. Despite widespread use, tractography has well-known pitfalls that limits the anatomical accuracy of this technique. Numerous modern methods have been developed to address these shortcomings through advances in acquisition, modeling, and computation. To test whether these advances improve tractography accuracy, we organized the ISBI 2018 3D Validation of Tractography with Experimental MRI (3D-VoTEM) challenge. We made available three unique independent tractography validation datasets – a physical phantom and two *ex vivo* brain specimens - resulting in 176 distinct submissions from 9 research groups. By comparing results over a wide range of fiber complexities and algorithmic strategies, this challenge provides a more comprehensive assessment of tractography’s inherent limitations than has been reported previously. The central results were consistent across all sub-challenges in that, despite advances in tractography methods, the anatomical accuracy of tractography has not dramatically improved in recent years. Taken together, our results independently confirm findings from decades of tractography validation studies, demonstrate inherent limitations in reconstructing white matter pathways using diffusion MRI data alone, and highlight the need for alternative or combinatorial strategies to accurately map the fiber pathways of the brain.

## Introduction

Mapping the detailed structural connectivity of the human brain has been a major scientific goal for decades. Currently, the only safe, non-invasive method to map the white matter connections in the living brain is called diffusion MRI tractography [1], which uses information about the displacement of water molecules in the brain [2] to map fiber pathways. For nearly two decades, tractography has been used to probe both the spatial extent (or trajectory) of white matter pathways, as well as the region-to-region (cortical-cortical) connectivity of the brain. These techniques have been applied not only by neuroscientists in order to elucidate fundamental insights about brain function, cognition, and development, as well as neurological diseases and disorders, but also by neurosurgeons for surgery planning [3]. Thus, the anatomical accuracy of tractography is critical for sound scientific conclusions or effective surgical outcomes. Specifically, tractography must be able to classify the presence or absence of connections in the brain (i.e. have high specificity and sensitivity), as well as precisely delineate the full spatial extent of the fiber pathways.

A number of validation studies have been carried out with the aim of determining the reliability of tractography - typically utilizing numerical simulations, physical phantoms, histological tracers, or comparisons against prior anatomical knowledge [4-23]. Together, this collection of studies have revealed pitfalls, uncertainties, and sources of error in the tractography process that may limit anatomical accuracy. For example, the sources of error can emerge during any stage of the tracking process: image acquisition, local voxel-wise reconstruction, and/or tracking streamlines from voxel to voxel. Specifically, with regard to image acquisition, it is well known that diffusion MRI (particularly with echo planar imaging (EPI)) is noisy, and prone to artifacts due to EPI susceptibility, head motion and eddy currents. These artifacts can lead to uncertainty in orientation estimates [4, 5], biases in diffusion indices [6], geometric distortion in pathways [7], all of which can result in anatomically incorrect tractography [8]. Another source of error involves drawing inferences about local fiber orientation from the diffusion displacement profile. MRI voxels are typically on the scale of millimeters, and can contain hundreds of thousands of axons with a large number of potentially complex geometric configurations (see [24] for a review on diffusion validation and its relationship to basic brain anatomy). In particular, fibers with crossing, kissing, fanning, and curving configurations have been a subject of concern for many diffusion reconstruction algorithms [9, 10, 23], resulting in incorrect and ambiguous estimates of fiber orientation [11]. In addition, these reconstructions have been shown to be dependent on data acquisition conditions (including signal-to-noise ratio, amount of diffusion weighting, and number of diffusion encoding directions), as well as axonal geometry (for example, the crossing fiber angle) [12, 13]. Finally, the tracking process itself is known to be subject to biases or inaccuracies due to lengths of streamlines [14], shape and size of pathways [15], cortical folding patterns [16, 17], ambiguity in pathways selection [18], and choices of tracking parameters (e.g., seeding and stopping criteria, step size, curvature thresholds) [19, 20]. Together, these difficulties have limited the anatomical accuracy of past tractography algorithms [21, 22]. Some authors even argued that the anatomical accuracy of diffusion MRI tractography is inherently limited because inferring fiber direction information from a water diffusion displacement profile is fundamentally a complex, underdetermined inverse problem [21].

Recently, several advancements in image acquisition, diffusion modeling, computational strategies, and tracking algorithms have been achieved with the aim of addressing these tractography limitations. To test whether these developments improve tracking accuracy, we organized the ISBI 2018 3-D Validation of Tractography with Experimental MRI (3D-VoTEM) challenge, which advances tractography validation using three different validation datasets: [1] a macaque dataset with a histological map of known tracer connections [21], [2] a squirrel monkey dataset with registered histological sections of the same sample, and [3] a physical fiber phantom with manually traced ground-truth pathways (Synaptive Medical, Toronto, ON).

This challenge differs from the conventional methods of validating tractography – rather than a researcher proposing a novel method or algorithm and evaluating this technique on proprietary datasets which can vary in a number of aspects, 3D-VoTEM provides image data and a reference standard to a number of independent research groups who can implement, parameterize, and optimize their choice of algorithms. Thus, this challenge serves as a platform to compare algorithms and results on the same data, and in a fair manner. Providing the community with three well-characterized, curated diffusion MRI and corresponding ground truth data, allows groups that may not have the resources or abilities to carry out animal experiments, histological processing, phantom construction, or MRI acquisitions to test their methodologies. In this way, this challenge facilitates validation from research groups that one group, acting alone, may be unable to perform due to limited resources, expertise, or hardware. In addition, tractography is performed by research groups that have tuned their setup for optimal performance, given their knowledge and experiences, rather than an individual research group evaluating many methods by simply evaluating an entire parameter space for one optimal solution of parameters. Finally, utilizing three independent sub-challenges allows us to test the conclusions that individual research groups have shown in the past. By evaluating the results of these three sub-challenges, each providing insights into the same problems, we sought to characterize the anatomical accuracy of the current state-of-the-art of diffusion tractography methods. In addition, by comparing results across a range of validation strategies, fiber complexities, and algorithmic strategies, the results from this challenge confirm the pitfalls of tractography revealed by independent research groups, as well as provide a more comprehensive assessment of tractography’s inherent limitations and successes than has been demonstrated previously.

## Materials and Methods

### Data and Ground Truth

The sub-challenges vary in both data acquisition and definition of ground truth. Example data and ground truth volumes are shown in Figure 1. The first sub-challenge consisted of a high quality - high resolution, high signal-to-noise ratio, and high angular sampling - ex vivo macaque dataset (Figure 1A) featured in previous validation studies [17, 21].

**Figure 1.**
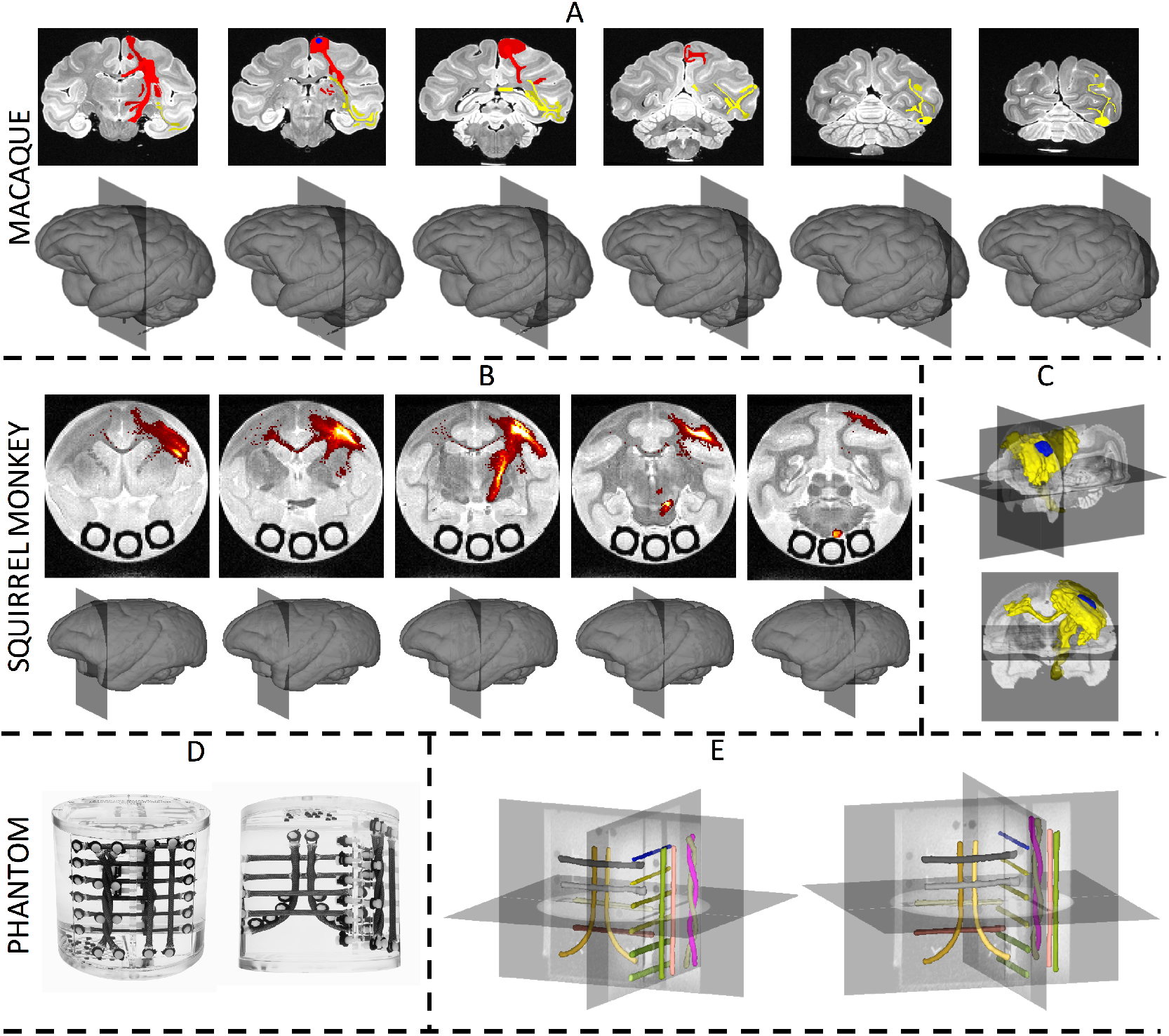
Ground truth fiber pathways for all sub-challenges. The macaque challenge ground truth (A) is derived from tracer studies for both the PCG (red labels) and V4v (yellow labels) pathways. The squirrel monkey ground truth is derived from histological tracer density maps from the same subject (B), and visualized as a 3D volume rendering (C). The phantom (D) ground truth for all bundles is derived from manual tracing on a high resolution T1 weighted image (E).

The two ground truth pathways were derived from anterograde tracer injections placed in the precentral gyrus (PCG) (Figure 1A, red) and the ventral part of visual area V4 (V4v) (Figure 1A, yellow), as described and characterized in [25]. Gray and white matter regions of interest were manually delineated on the data in order to assess agreement between tracer and tractography. This dataset allows validation of region-to-region connectivity. The second sub-challenge is performed on an ex vivo squirrel monkey dataset [26, 27], acquired at a coarser resolution (relative to brain volumes), a lower SNR, and fewer sampling directions (31 versus 114 for the macaque). The ground truth, is defined based on an anterograde and retrograde tracer injection in the primary motor cortex (M1) of the same brain. Image processing on histological slices allows extraction of the ground truth fiber pathways on a voxel-by-voxel basis (Figure 1B) as well as the creation of a binary “ground-truth” fiber pathway (Figure 1C). Gray and white matter regions of interest are defined based on additional histological stains. This subchallenge allows validation of both region-to-region connectivity as well as voxel-wise spatial overlap between tractogram (tractography streamlines) and tracer. The final subchallenge consists of data acquired on a biomimetic anisotropic diffusion phantom (Synaptive Medical, Toronto, ON) containing 16 separate fiber bundles (Figure 1D). Image acquisition consists of an overnight scan on two different scanners (scanner “A” and scanner “B”) in the same imaging facility, with multiple diffusion weightings (b=1,000 and 2,000 s/mm2), a large number of sampling directions (96 per b-value), and seven repetitions. The ground truth is manually defined on a high resolution T1-weighted image for all 16 bundles, and registered to dMRI space for a voxel-wise comparison of the spatial overlap between tractography and ground truth bundles (Figure 1E). Details regarding the acquisition and processing procedures, as well as ground truth creation, are described below.

### Sub-Challenge 1 - Ex Vivo Macaque

#### Data Description

The provided dataset is the one used and described in detail in [21]. Briefly, the images were acquired from an ex-vivo fixed macaque brain at 0.25mm isotropic resolution. The diffusion weighted-images (DWIs) contain 7 volumes with b=0 s/mm^2^ and 114 volumes with b=4900 s/mm^2^ (with small variations due to the effects of the imaging gradients). The data were preprocessed using the TORTOISE software package [28] and were corrected for eddy current distortions and motion-like artifacts caused by frequency drifts.

#### Ground Truth Pathways

Two ground truth pathways were derived from the anterograde tracer injections placed in (i) the precentral gyrus corresponding to the foot region of the primary motor cortex and (ii) rostroventral part of the occipital region corresponding to the ventral part of area V4 (V4v) and the adjacent ventral area V3 – as described and characterized in [25]. The tracer-labeled regions of interest were transferred to the same space as the diffusion data. In addition, gray matter and white matter regions of interest were manually delineated on the high-resolution data in order to assess agreement between tracer and tractography results.

### Sub-Challenge 2 - Ex Vivo Squirrel Monkey

#### Tracer Injection

The histological ground truth data is acquired on a squirrel monkey brain. Here we utilize a commonly used neuroanatomical tracer for studying neuronal pathways, biotinylated dextrane amine (BDA). Because it is transported both anterograde and retrograde, BDA yields sensitive and detailed labeling of both axons and terminals, as well as neuronal cell bodies. This tracer relies on axonal transport systems; thus BDA injection is performed prior to ex vivo imaging. Under general anesthesia using aseptic techniques, BDA was injected into left hemisphere M1 cortex. Eight injections were made in order to cover a large M1 region representing the forearm as identified by intracortical microstimulation. After surgery, the monkey was allowed to recover from the procedure, giving the tracer sufficient time to be transported along axons to all regions connected to M1.

#### MRI Imaging

For ex vivo scanning, the brain was perfusion fixed with 4% paraformaldehyde preceded by rinse with physiological saline. The brain was removed from the skull and stored in buffered saline overnight. The next day, the brain was scanned on a 9.4 Tesla Varian scanner. Diffusion-weighted imaging was performed using a pulsed gradient spin echo multi-shot spinwarp imaging sequence with full brain coverage (TR=5.2s, TE=26 ms, number of diffusion gradient directions=31, b=0, 1200s/mm2, voxel size=300×300× 300 μm3, data matrix=128×128×192, number of acquisitions=10, SNR«25, scanning time«50 hr). The b value used in this experiment was lower than is optimal for diffusion studies in fixed tissue, due to hardware limitations. A low b value decreases the available diffusion contrast-to-noise ratio (CNR) in the image data, which has the same effect as higher image noise. To compensate for this shortcoming, we extended the scan time to 50 hours, which yielded a CNR comparable to in vivo human studies (equivalent to an in vivo study with mean diffusivity=0.7×10-3 mm2/s and SNR«20). All sub-challenge data will be distributed and analyzed directly in the space in which diffusion data were acquired.

#### Histological Acquisition

Following ex vivo MRI scanning, the brain was frozen and cut serially on a microtome in the coronal plane at 50 um thickness. Prior to cutting every third section (i.e., at 150 mm intervals), the surface of the frozen tissue block was photographed using a Canon digital camera (image resolution = 50 um/pixel, image size = 3330×4000 pixels, number of images per brain ~ 280), mounted above the microtome. Every 6th section (approximately the size of an MR voxel) is processed for BDA to trace pathways associated with the M1 cortex. Whole-slide Brightfield microscopy was performed using a Leica SCN400 Slide Scanner at 20x magnification, resulting in a maximum in-plane resolution of 0.5um/pixel.

#### Ground Truth M1 Connectivity

The “ground truth” connectivity of the injection area was determined by the presence of BDA-labelled axons in our high-resolution histology, which displayed as brown in the digital images. BDA-labeled fibers were segmented and counted following a series of morphological processes: top-hat filtering was performed to correct uneven illumination, global thresholding to extract fibers (segmenting brown [r/g/b = 165/42/42] using the “colorseg” function available on MathWorks File Exchange), and morphological operations to remove non-fiber objects (objects less than 11 pixels, empirically chosen) and to remove branch points of overlapping fibers. Histological images were downsampled to the resolution of the MRI-data (300 um isotropic), and the number of BDA fibers per voxel was counted, resulting in BDA density maps. These BDA density maps represent the ground truth “strength of connections” to the M1 injection area.

A total of 71 gray matter and white matter regions of interest were defined in MRI-space, using both histological and MRI-derived information, as described in [26, 29, 30], and retrieved from the squirrel monkey brain atlas [27], in order to assess connectivity agreement between tracer and tractography.

#### Registration

The multi-step registration utilized here is very similar to the registration procedure validated in an earlier study [31], which showed that the accuracy of the overall registration was approximately one MRI voxel (~0.3 mm). From the Leica image file, the TIFF image stored at 128 um/pixel (down-sample factor 256) was extracted and registered to the down-sampled photograph (256×256 pixels at a resolution of approximately 128 um/pixel) of the corresponding tissue block using a 2D affine transformation followed by a 2D non-rigid transformation, semi-automatically calculated via the Thin-Plate Spline algorithm [32]. Next, all down-sampled block face photographs were assembled into a 3D block volume and registered to the corresponding 3D MRI volume using a 3D affine transformation followed by a non-rigid transformation automatically calculated via the Adaptive Bases Algorithm [33]. The deformation fields produced by all registration steps were applied to processed histological data in order to transfer the ground truth histological pathways into the diffusion space for comparisons with tractography.

### Sub-Challenge 3 - Anisotropic Fiber Phantom

#### Phantom Construction

The Anisotropic Diffusion Phantom is a biomimetic phantom containing complex geometries of anisotropic fibers that mimic the tissues of the brain. The phantom contains 16 flexible fiber bundles. Pathways are aligned in orthogonal planes, as well as in curved (both 90 degrees and helical curving), and kissing geometries (as well as a resolution bank) to mimic complex nerve fibers of the brain, with bundle dimensions of magnitudes comparable to major white matter pathways in the human brain.

#### MRI Imaging

MR scans were performed on two scanners, both Philips 3.0T systems. The 16 cm diameter phantom (matrix fluid filled) was imaged for both structural and diffusion contrasts. The structural scan utilized a 3D MPRAGE sequence to acquire a T1 contrast (TE/TR = 3.6/8ms, Matrix = 256 * 256, Resolution = 0.88*0.88mm, slice thickness = 1. 0mm). A low-resolution diffusion contrast was acquired using a 2D EPI diffusion weighted sequence (TE/TR = 75ms/9.65s, Matrix = 72*72, resolution = 2.25*2.25mm, slice thickness = 2.5mm). 96 diffusion directions were acquired, uniformly sampled over a sphere, at b-values of 1,000 s/mm2 and 2,000 s/mm2. Non-diffusion weighted images were acquired between every 8 diffusion-weighted images. Sampling was performed with phase encoding both anterior to posterior, and repeated posterior to anterior, in order to allow pre-processing for motion, eddy currents, and susceptibility distortions. This series of scans (2 b-values, 96 uniformly distributed directions, with two phase encoding directions each) was repeated 7 times on each scanner.

#### MRI data processing

Diffusion MRI pre-processing was performed in the coordinate system that the data were acquired in. Steps included correction for movement, susceptibility induced distortions, and eddy currents using FSLs *topup* and *eddy* algorithms [5]. The gradient tables were rotated based on the transformations obtained from the corrections.

#### Ground Truth

Ground Truth was manually delineated for each bundle on the T1-weighted high-resolution image, separately for each scanner, using ITK Snap (www.itksnap.org, v2.4.0). For each scanner, the T1-weighted image was registered to the average nondiffusion weighted image using 3D affine followed by a 3D non-rigid registration (FSL Software Library v5.0 [34]). Ground truth labels were individually transformed to diffusion space using nearest-neighbor interpolation.

### Anatomical Accuracy Measures

Measures were calculated which describe the anatomical fidelity of the resulting tractograms, several of which have been previously employed in the validation literature. Here, measures are divided into voxel-wise fidelity metrics, and ROI-based fidelity metrics. In the following, the Ground Truth volume is represented by G_j_ (j = 1,2, …, m) and tractography volume represented by T_i_ (i=1,2, …, n).

#### Voxel-wise measures

- Bundle Overlap (OL) [22]: The proportion of voxels that contain the ground truth volume that are traversed by at least one streamline. The OL describes how well tractography is able to describe the volume occupied by the ground truth and is defined as:

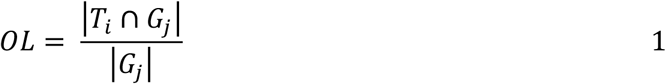

where | • | denotes cardinality.
- Bundle Overreach (OR) [22]: the number of voxels containing streamlines that are outside of the ground truth volume divided by the total number of voxels within the ground truth bundle:

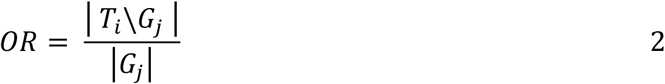

where operator \ denotes relative complement operation.

All voxel-wise measures were calculated for the phantom and squirrel monkey subchallenges, because the ground truth volumes are defined voxel-wise volume.

#### ROI-based measures

For both squirrel monkey and macaque sub-challenges, the ROI-based connectivity to seed regions was assessed using the white matter and gray matter regions of interest. Anatomical fidelity metrics of sensitivity, specificity, and Youden index were derived for all tractograms.

- Sensitivity – True positive rate; measures the proportion of positives (regions that are occupied by ground truth) that are correctly identified as such (using tractography). Sensitivity measures the ability to correctly detect all connections of the seed region.
- Specificity – True negative rate; measures the proportion of negatives (regions that do not contain ground truth) that are correctly identified as such (do not contain streamlines). Specificity measures the ability to correctly identify voxels that do not have connections with the seed region.
- Youden’s J statistic – Sensitivity+Specificity-1; a statistic that captures the performance of a diagnostic test, and estimates the probability of an informed decision, ranging from -1 to 1. A value of 1 indicates a perfect test with no false positives or false negatives.

All metrics, both voxel-wise and ROI-based are computed for all algorithms.

## Results

### Submissions

Although the submission site remains open (https://my.vanderbilt.edu/votem/submissions/), the data in this study includes only those submitted before the ISBI 2018 conference (April 4, 2018). Overall, 176 unique submissions were submitted across the challenges (58 for the macaque, 62 for squirrel monkey, and 56 for the phantom) from nine international research groups. Submissions ranged in complexity from open-sourced software, diffusion tensor based tractography with default software configurations to that of complex, multi-shell, in-house algorithms with extensive post-processing – with most featuring either reconstruction or tracking strategies developed in the last few years. Details of each submission are provided in Supplementary Tables 1-3. The most common reconstruction methods were some form of spherical deconvolution or multi-compartment models. Both deterministic and probabilistic algorithms were employed, with most utilizing some form of constraint on fractional anisotropy (FA), curvature, or anatomical mask. The seed regions (where tractography is initiated) provided along with the datasets were used as both true seeds as well as regions of interest after whole-brain tractography was performed. Standard pre-processing for susceptibility distortions, motion, and eddy currents was performed for all datasets, but very few groups used additional pre-processing steps (with the exception of denoising techniques), and post-processing included various filtering techniques, track grouping, and manual track selection.

### Qualitative Results

The tractography streamlines for randomly selected submissions are shown in Figure 2 for the three sub-challenges. Qualitatively, there is large variability in the resulting connectivity profiles and pathways represented. Specifically, for the macaque and squirrel monkey, visualizing submitted streamlines shows a range in spatial extent from only connectivity nearby the seed region, to covering large expanses of the entire hemisphere. The phantom submissions generally capture the correct shape, position, and orientation of all 16 bundles, with most noticeable differences in sparsity of streamlines and thickness of pathways.

**Figure 2.**
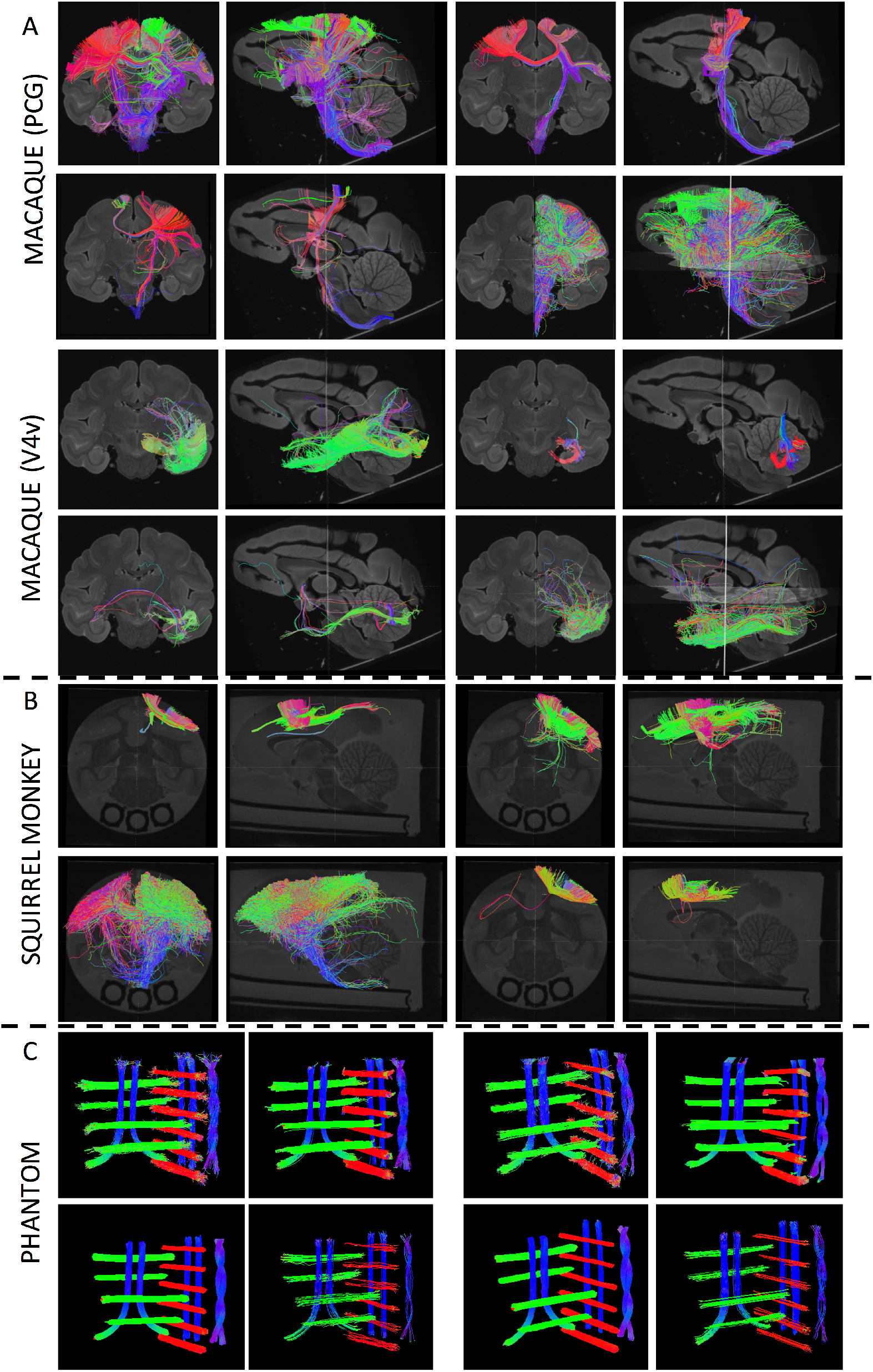
Diffusion tractograms for randomly selected submissions. Tractography is shown in the coronal and sagittal planes, for both macaque pathways (A), the squirrel monkey pathway (B), and all 16 phantom bundles (C).

### Region-to-region connectivity validation – Sensitivity and Specificity

For the macaque and squirrel monkey datasets the agreement between tracer and tractography results are evaluated using sensitivity and specificity measures, validating the ability of tractography to accurately map region-to-region (or seed-to-region, see Materials and Methods section) connectivity. Additionally, to identify the best combination of sensitivity and specificity, the Youden index (J) (Specificity + Sensitivity – 1) is computed, where a value of 1 indicates a perfect test and a value of zero indicates no predictive value. The results across all submissions are shown as ROC curves in Figure 3, where marker color indicates unique research groups. In both macaque and the squirrel monkey datasets, the main finding is that no algorithm or submission consistently identifies true positive pathways without also generating a large number of false positive pathways, and none consistently identify true non-connections without suffering a low true positive rate (i.e., an increase in sensitivity comes at the cost of a decrease in specificity, and vice-versa). For the macaque, most submissions result in high specificity values (with a large number of false negative connections), while the squirrel monkey algorithms typically lie at the extremes of the ROC plots.

**Figure 3.**
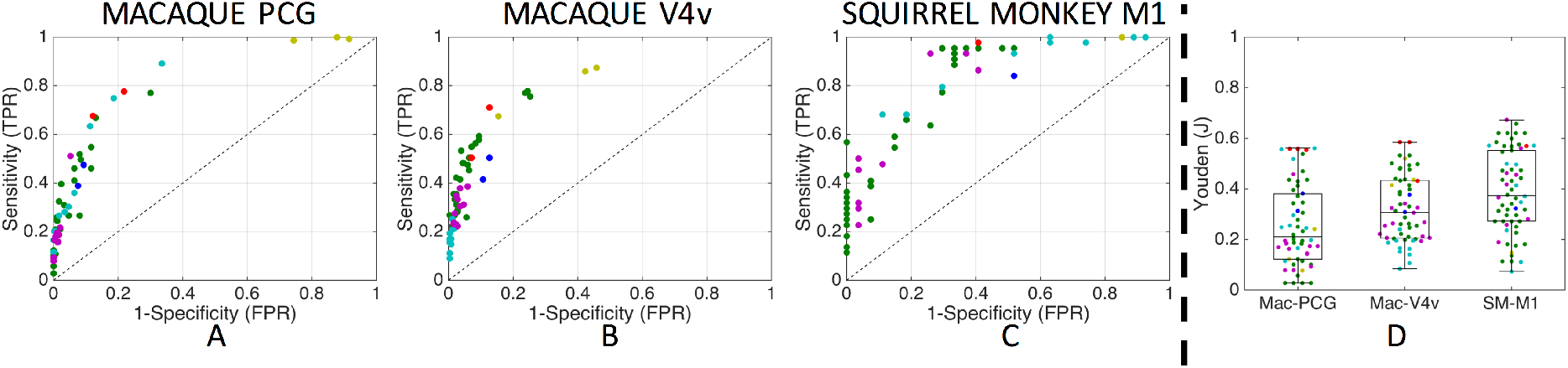
Region-to-region connectivity validation. Results are shown as ROC curves on the macaque PCG (A) and V4v (B) pathways, and squirrel monkey M1 pathways (C). Boxplots of corresponding Youden values (D) are shown for both challenges. One marker is shown per submission, with marker colors indicating unique research groups.

Most submissions have relatively low predictive value, with median Youden indices of 0. 21, 0.30, and 0.37 for macaque PCG, macaque V4v, and squirrel monkey M1 pathways, respectively (Figure 3D). The highest Youden values for each pathway are only 0.56, 0.58, and 0.67. Thus, even the anatomical accuracy of the most predictive algorithms are suboptimal. The squirrel monkey results have a statistically significant (1way ANOVA, p<0.01) higher population mean Youden value than the macaque results - thus, in general, the algorithms provide slightly more anatomically accurate tracts on the squirrel monkey than macaque.

### Spatial Overlap validation – Bundle Overlap and Overreach

A voxel-wise measure of spatial agreement between tracer and tractography is possible for the squirrel monkey with binary tracer data and phantom datasets with manually drawn tracts, because the ground truths are established in the same animal/phantom, making voxel-by-voxel comparisons possible. In these sub-challenges, we compute the bundle overlap: the proportion of voxels that contain the ground truth that are traversed by a streamline – and bundle overreach: the number of voxels containing streamlines outside the ground truth divided by the total number of voxels within the ground truth. In short, the overlap is a measure of the true positive rate (i.e., sensitivity) while the overreach is related to the false positive rate (i.e., specificity).

Plots of overlap and overreach for the squirrel monkey and both phantom scans (Figure 4, A-C) show very similar results as the regional connectivity accuracy: algorithms that are successful at identifying the full extent of the pathways (high overlap) suffer from high overreach. In the squirrel monkey, algorithms that did not suffer from a significant overreach (<10%), often had very low overlap values, identifying less than 25% of the full histologically defined ground truth volume. While the phantom had significantly improved overlap values, many algorithms that recover the full bundle volumes can suffer from overreach as much as 1.5-5x the actual ground truth volume.

**Figure 4.**
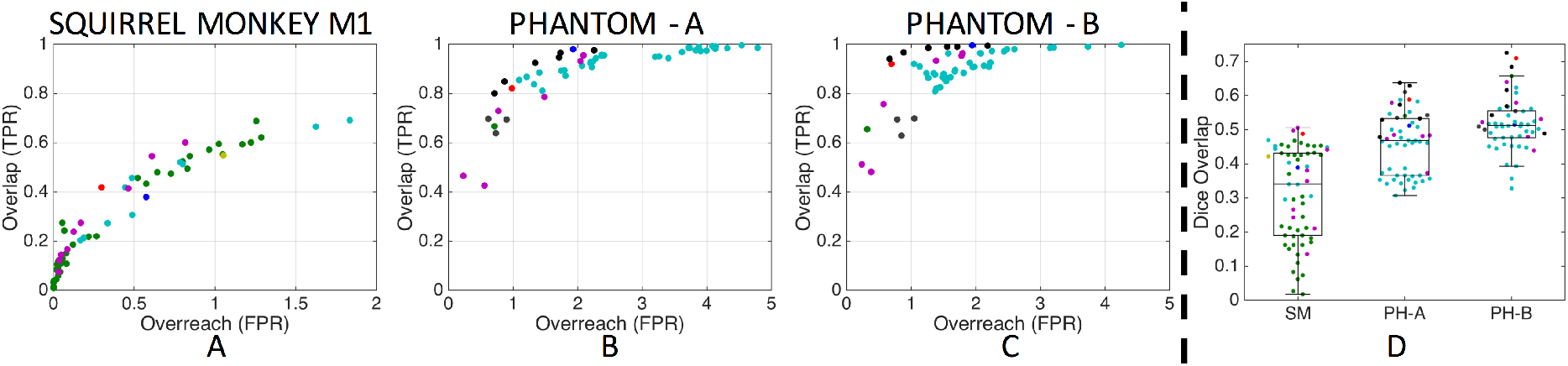
Voxel-wise spatial overlap validation. Plots of overlap versus overreach are shown for the squirrel monkey (A) and phantom datasets on scanner A (C) and scanner B (C). Boxplots of the corresponding Dice overlap coefficients are shown for both challenges. One marker is shown per submission, with marker colors indicating unique research groups.

The Dice overlap coefficient (Figure 4, D) has median values of 0.34, 0.46, and 0.51 for the squirrel monkey, phantom on scanner A, and phantom on scanner B, respectively, with maximum Dice coefficients reach 0.51, 0.63, and 0.72. The phantom submissions have statistically significant (1-way ANOVA, p<0.01) higher Dice coefficients than that of the squirrel monkey, indicating an overall better voxel-wise accuracy.

## Discussion

The 3D-VoTEM challenge combines and presents three separate tractography validation strategies, inviting ideas and algorithms from researchers from around the world, with the primary objective to determine whether recent technical advancements in diffusion MRI tractography can deliver anatomically accurate maps of the brain structural connectivity. More specifically, given the known limitations of these techniques, we asked if advances in algorithms, acquisition, and methodologies utilized in modern tractography techniques have improved anatomical accuracy. The key finding is that, despite a better understanding of limitations and pitfalls of these techniques, and considerable effort leading to advances in these algorithms, the anatomical accuracy of modern tractography approaches is still limited. Importantly, the limited anatomical accuracy is observed in three independent sub-challenges, each with algorithms created, developed, and optimized by leading research groups in the field. These findings support the results and conclusions demonstrated over the last decade of validation studies, across species and phantoms, performed by individual research groups.

## Limits to Accuracy

Importantly, we find consistent results across a diverse range of validation approaches. The sub-challenges vary in not only the systems under investigation (phantom versus non-human primates), but also acquisition (voxel size, angular resolution, SNR, diffusion weightings), complexity of pathways, and definition of ground truth. In all cases, algorithms that succeeded in recovering the true connections (high sensitivity or high overlap) consistently generated a large number of false positive connections (low specificity or high overreach), and no algorithm was highly informative or highly similar to the ground truth (high Youden or high Dice). In fact, most algorithms had surprisingly low connectivity predictive value and low spatial overlap with the true pathways. Thus, accuracy in tractography is not only hampered by a false positive problem [18], but many algorithms appear to be dominated by false negative connections.

While the accuracy tradeoffs have been consistent across challenges, differences in tractography performance between the challenges are apparent. This is expected, as tractography, and especially local reconstruction, are known to be heavily affected by the quality of the diffusion MRI acquisition. For example, it is generally assumed that many failures of tractography will be mitigated through improved angular and spatial resolution data. However, tractography in the macaque system resulted in less accurate connectivity measurements than tractography in the squirrel monkey system, despite significantly improved resolution, SNR, and diffusion sensitivity. Thus, differences in accuracy likely depend on the complexity of the pathway of interest, rather than acquisition quality alone. It should also be considered, however, that the ground truth data for the macaque brain were obtained from tracer studies performed in different animals so that interindividual variability in brain connections may have slightly lowered the accuracy value that could be reached with that dataset. Similarly, the phantom, with relatively sparse, well-defined, and less-complicated pathways resulted in significantly higher overlap agreement than that of the squirrel monkey.

We consider all submissions, in all challenges, to be “modern” algorithms. In most cases, investigators implemented reconstruction or tractography techniques developed only recently, with many specifically created to address one or more known limitations. Most importantly, these algorithms were tuned based on the collective knowledge and experience of the research lab, with the aim to optimize the accuracy of their results. Other implemented algorithms were proposed as early as 2001 (for example, using the tensor with a low-order streamline integration), and while they may be considered rudimentary or basic, because they are still in use today – sometimes as the default algorithm in many open source software packages – they are considered modern. Thus, the observed plateau or limits in anatomical accuracy applies to not only the state of the art approaches, but also to the techniques of the past, on which the bulk of current knowledge of structural connectivity in the human brain is based upon.

The trend in many of the more recently developed algorithms and pipelines is to include some variation of informed post-processing. This includes track grouping or clustering [35], streamline filtering based on the diffusion signal or track densities [36], globally fitting streamlines to microstructural models [37], and even manual delineation of regions of interest or streamlines. In all 67 submissions (33 phantom, 11 squirrel monkey, 23 macaque) used some form of either anatomically-, globally-, or microstructurally-informed post-processing. Although the false positive rate was reduced in many of these (increased specificity, decreased overreach), no statistically significant difference was observed between these and submissions not utilizing postprocessing – although there is a diverse range of alternative confounding factors across algorithms, including pre-processing, reconstruction methods, algorithms, constraints and number of streamlines. It would be informative to compare tractograms to ground truth both before and after post-processing to confirm increased accuracy and reduced false positives. In addition to new post-processing, several teams used recently developed reconstruction methods (most a variant of spherical deconvolution or multicompartment models), software packages (Dipy, MI-BRAIN, Quantitative Imaging Toolkit, dMRITool, MRTrix, FiberNavigator), and streamline algorithms.

The results of the 3D VoTEM challenge confirm and expand upon the limitations and shortcomings demonstrated over the last decade in validation literature. Importantly, the algorithms submitted in this challenge are run and optimized (and often developed) by the contestants themselves, rather than run as off-the shelf algorithms typically implemented in validation literature. Submitted algorithms are compared and benchmarked on the same dataset, using the same evaluation criteria. In the past, both white matter pathways and long-range connections have been assessed using either histological validations [14, 19, 38-40], simulated datasets [41], or physical phantoms [22]. Past studies have demonstrated that DTI tractography has difficulties when tracts cross or divide [38], highlighting the importance of the crossing fiber problem. However, DTI tractography is strongly correlated with true connectivity on the scale of major cortical regions, but is less reliable at measuring voxel-wise connectivity [42]. The current challenge confirms this, not only for DTI, but for a range of reconstruction techniques (Both DTI and higher order models) and tracking strategies. Cortical-cortical connection strengths of tractography have been shown to be modestly informative predictions of tracer connections [14] with biases dependent on path lengths and connections strengths. Tractography is also capable of finding the spatial extent of major pathways [19], however, it was found not possible to achieve high specificity and sensitivity at the same time, with only moderate ability to detect true positive (~0.35-0.85 true positive rate) and true negative (~0.05-0.4 true negative rate) connections. General conclusions across all studies were that tractography was informative, but that accuracy would be improved through improvements in acquisition, newer algorithms, high quality data. Towards this end, in 2013, Thomas et al. [21] acquired an ex vivo macaque dataset with high angular and spatial resolution – estimated to be equivalent to an in vivo acquisition requiring thousands of hours of scan time. Using standard algorithms at the time, they find that despite exceptional data, accurate tractography still remains an elusive goal. In comparison, even with new and improved algorithms in the current macaque sub-challenge, only minor improvements are made (an increase in Youden value of 0.05 for the optimal algorithm) in accuracy compared to those from nearly five years ago – suggesting that the ROC curves have not shifted dramatically in the last few years.

## Advancements Needed

While our phantom and ex vivo validations result in similar trends and findings across a range of ground truth geometries, acquisition settings, and image qualities, the ultimate goal is to accurately map the *in vivo* human brain. Although tractography on a human cannot be directly validated, the accuracy of tractography based on these non-human validation paradigms has largely plateaued in recent years, which likely reflects similar sensitivity/specificity limitations of the process in a human brain. These specific datasets should require a dedicated processing pipeline, tuned and optimized for it. Most existing tools and software packages were developed and tuned based on the field’s understanding of human anatomy. While some exist (or are easily adaptable) (https://osf.io/yp4qg/), tools for small animals, or larger animals e.g. monkeys, or any non-human tractography need to be improved to create better masks, better labels, or better priors so that modern, and future, tractography developments can be leveraged. These tools and resources will not only be applicable to validation studies, but any research on the structural connectivity of the non-human brain.

“Solving” the tractography problems in these phantoms and animal models does not necessarily guarantee perfect reconstructions in the human brain. However, better understanding mistakes relative to the ground truth will certainly spur improvements and innovations in these techniques. Advances will be made through a process well-described by Dyrby et al. [24] where we must “loop until our method’s results agree with the gold standard, and/or until the updated knowledge of ground truth can explain the discrepancies observed.” This includes continually updating theory and implementation of methods, validation against gold standards, understanding deviations from the truth, followed by further modifications to theory and implementation, etc. Consequently, there is a need for more advanced and sophisticated gold standards, and a need for validation across a range of spatial scales. In the past (and in the current study), validation is done as an overall assessment in sensitivity and specificity (or overlap and overreach). Future studies should not only explore accuracy at assessing connections and overlap, but also voxel-wise and microstructural features of the datasets. For example, validation strategies could include multiple histological stains or phantoms with varying fiber densities/diameters/volume fractions, in order to evaluate both connectivity and microstructural features simultaneously. The multi-modal or multiscale strategies could lend insight into individual steps of the tracking process in order to better understand where tractography “first” goes wrong – whether it is assumptions about microstructural features, axonal orientations, or simply tractography decision making.

When validating tractography it is important to clearly define what we hope to map with tractography, and more importantly, how well the ground truth represents this. The goal could be to validate microstructural features of specific pathways (fiber densities, fiber orientations), the course of white matter pathways, the presence or absence of connections between regions, or some measure of connectivity between regions (number of connecting axons, proportion of axons reaching a region, conductivity between regions). While the challenges in this study focused on the course of the pathway (phantom and squirrel monkey) and presence or absence of connections (macaque and squirrel monkey), they are not without their limitations in representing true tissue structures [24]. Several factors limit the accuracy of the gold standard in ex vivo validations, including changes in tissue due to extraction and fixation, imperfect registrations between histology and MRI, and tracer uptake and visualization. As mentioned above, the macaque MRI and tracer injection was performed on different subjects. While the squirrel monkey experiments were all on the same subject, the acquisition was sub-optimal for ex vivo imaging [43], and included only a single pathway of interest. The phantom is limited by its simplicity, with a simple geometry on the macroscopic scale. Potential opportunities involve including more adjacent bundles (crossing, kissing, fanning) where partial volume occurs on the scale of individual voxels, as well as features that better mimic the in vivo brain (cortex, varying diffusion compartments, fiber dispersion). Future validation approaches should continually strive for improvements in creation or construction of the ground truth, aim for innovation in validation approaches and strategies, and aim to minimize deviations of the “ground truth” from the true tissue properties by accurately extracting the feature of interest.

This stresses the need for sharing and distribution of validation datasets and ground truths, and tackling the validation problem from a number of perspectives is critical. However, these datasets are time consuming to acquire, expensive, and often require expertise in various niche fields (i.e. histology or phantom creation). While the current challenge was the first to combine separate datasets with very different validation strategies, there are a large number of existing datasets that have lent their own, unique, insight into interpreting tractography (see above for examples). However, it is important to not only validate tractography on different spatial scales (i.e. microscopic versus macroscopic), diverse datasets, and various representations of ground truth, but also necessary to make these open-source for valid comparisons of existing and future algorithms and approaches. An online tractography validation tool (much like the “Tractometer” tool for the FiberCup physical phantom [22]) combining a large repository of validation datasets would make it easier for neuroscientists, computer scientists, and physicians to submit and test new algorithms, datasets, and methods. Current neuroimaging validation databases do exist (https://osf.io/yp4qg/), containing largely microstructural validation datasets – but tractography is just modeling microstructure at a macroscopic length scale. Thus, we recommend this, or similar, databases to collect and distribute tractography validation data. This, in combination with more sophisticated algorithms, will almost certainly lead to advances in tractography, and allow us to gain better insights into trends and limitations of these techniques.

While it seems that the results of this study paint a pessimistic view of tractography, there are several positive takeaways. First, some algorithms are indeed able to recover the full spatial extent of pathways, while others have a specificity high enough to make confident predictions about the presence of pathways. Finally, reassuringly, there will almost always be human involvement in this process, especially if tractography is used for surgical planning. A surgeon may not be interested in sparse, stray tracts, or may only care about streamlines in specific locations (i.e. peri-tumoral), and perfect sensitivity/specificity may not be a concern. Alternatively, interaction with the tracking software (and subsequent parameters, ROIs, etc.) allows the surgeon to fine tune based on his or her prior knowledge. This, in combination with the large variability in reconstructions, makes it critical to educate tractography users that the process as it stands is more akin to an art, than an absolute representation of the brains fiber pathways.

In a typical use of tractography, an investigator uses estimated orientation information to ask which brain region is connected to another, as well as the shape, size, route, and strength of this connection. Similarly, in these challenges, the only information given to the investigator is in the form of fiber orientation information (the diffusion signal), and the beginning of the pathway (the seed region). Results from our current study as well as the seminal work of Maier-Hein et al. [18] clearly shows that having ***only this information, i.e. the local orientation and seed, is not enough!*** Tractography needs more information to overcome the specificity-sensitivity curse of current methods.

Potential solutions are appearing such as i) including better and more priors based on known neuroanatomy, ii) including microstructural information along local orientations to better trace-out orientations that belong to the same connection from end-to-end, iii) machine learning techniques that could learn from all submissions, from challenges with ground truth, the local and global structure of valid and invalid connections, and iv) information from other modalities such as myelin markers and functional imaging contrasts that could help reduce the number of invalid connection and increase the number of valid connections.

Better priors from hundreds of years of neuroanatomy research as well as functional imaging could bring novel information about the ‘where’ and ‘how’ streamlines should start and end, as well as traverse complex crossing and bottleneck regions. Microstructural information from dMRI or other modalities could add a vector of features along each fiber orientation to help connect orientations that belong to the same structure, that have the same properties (axon diameter, intra/extra-cellular space, myelin volume, etc). Moreover, with the terabytes of streamlines generated by state-of-the-art techniques in numerous challenges organized internationally as well as initiatives such as Tractometer, there is a great potential for having a deep learning algorithm learn the easy-to-track and hard-to-track parts of the brain, both locally and globally, and potentially highlight the untrackable regions and locations of errors.

While no submission was consistently successful in every tracking fidelity metric, the results of our study do not invalidate tractography as a useful biomedical tool, as many were fairly predictive of connectivity, or had moderate to good ability to delineate spatial pathways. Instead, the results of our study emphasize that given current state of the art approaches, pathway reconstruction increasingly appears to be a problem that is unlikely to be wholly solved using only local orientation estimates, and it may be necessary to incorporate other information, other modalities, or new tracking strategies, to successfully resolve tractography’s known limitations.

## Acknowledgments

This work was conducted in part using the resources of the Advanced Computing Center for Research and Education at Vanderbilt University, Nashville, TN. We thank Synaptive Medical for providing the anisotropic diffusion phantom, and for expertise on its use in research. This work was supported by the National Institutes of Health under award numbers R01EB017230 (Landman), and T32EB001628. This project was supported in part by VISE/VICTR VR3029 and the National Center for Research Resources, Grant UL1 RR024975-01. All data, for all three challenges, are freely available at the challenge website (https://my.vanderbilt.edu/votem/). Challenge submissions remain open, and will be processed upon submission.

## Supplementary Materials

See Supplementary Materials for tables describing details of each submission in detail, for each sub-challenge.

